# Positive and negative cognitive biases in a cockroach

**DOI:** 10.64898/2025.12.01.691180

**Authors:** Adam M. Guilfoyle, David N. Fisher

## Abstract

Understanding the mental experiences of animals is key for managing them and mapping the evolution of cognitive complexity. Emotional and affective states are fundamental to these goals as their presence indicates the potential for negative welfare states and so suffering. However, until recently the potential for affective states in invertebrates has been underappreciated, especially for positive affective states where research is limited to a single order. Here, we adapt cognitive bias tests to examine both negative (pessimistic) and positive (optimistic) bias in cockroaches (*Blaptica dubia*) in order to expand the taxonomic reach of affective state research and to provide a rare test for positive affective states in an invertebrate. We trained cockroaches to associate two different odours with either positive reinforcement or positive punishment and then assessed their reactions to ambiguous stimuli (a range of mixtures of the trained odours). We found that cockroaches exposed to light were more pessimistic (negative affective state) than controls, while those exposed to the scent of opposite sex conspecifics were marginally more optimistic (positive affective state) than controls. Our results are therefore consistent with the potential for affective states in cockroaches and for positive affective states in invertebrates more widely, indicating a previously unexpected level of motivational complexity.

## Introduction

An animal’s welfare depends on both its physical and mental wellbeing (Broom, 1991; Broom, 2016; Duncan, 2016). When assessing the mood or emotional state of humans, we can simply ask them how they feel. Assessing mental states akin to moods and emotions in non-human animals is more challenging as animals cannot verbally self-report and communicate their state of mind, well-being, and mood to researchers (Dawkins, 1990; Izard, 2009). Researchers typically rely on three approaches for studying emotional states in animals (Mendl *et al*., 2010b; Mendl *et al*., 2011; Bateson *et al*., 2011; Perry & Baciadonna, 2017): behavioural (e.g. fight or flight behaviours), physiological (e.g. hormone analysis), and cognitive (e.g. perception-based biases). Of the three, cognitive measurements (and their various bias tests) have become a particularly popular methodology for measuring moods and emotion-like states relevant to animal welfare (Mendl *et al*., 2009; Schuck-Paim & Alonso, 2023). Testing for cognitive bias involves testing for biases in the information processing of individuals, with biases towards negative stimuli indicating negative affective states, and vice versa (Lagisz *et al*., 2020).

The methods of studying the cognitive influences of affective states can be further split into three categories (Perry & Baciadonna, 2017): memory bias (e.g. memory retrieval), attentional bias (e.g. vigilance), and judgemental bias (e.g. risk-taking). Judgement bias testing has become a popular method of empirically measuring the presence and influence of affective states in animals and is often referred to as a proxy indicator of emotion (Mendle *et al*., 2009; Paul *et al*., 2011; Bateson *et al*., 2011; Godfrey-Smith, 2021b). Judgement bias testing measures decision-making biases to ambiguous stimuli under varying conditions, indicating an affective state’s presence (Bethell, 2015). A negative affective state (such as anxiety or stress) may lead to decreased interaction with ambiguous stimuli. These pessimistic cognitive biases have been reliably correlated with negative feelings in humans, have been regularly observed and studied in animals, and have been used as a correlating indicator of emotional (or emotion-like) states (Mendl *et al*., 2009; Paul *et al*., 2011; Bateson *et al*., 2011; Godfrey-Smith, 2021b). Consequently, judgement bias tests have become an increasingly common research method in animal welfare science and applied to various species of mostly captive vertebrates (Bethell, 2015; Lagisz *et al*., 2020): farm- and domesticated animals (Mendl *et al*., 2010a; Baciadonna & McElligott, 2023), terrestrial mammals (Bateson & Nettle, 2015; Hales *et al*., 2021), marine mammals (Clegg & Delfour, 2018; Delfour & Aviva, 2021), birds (Horváth *et al*., 2016; McCoy *et al*., 2019), reptiles (Benn *et al*., 2019), and fish (Langérôme *et al*., 2024; Brown, 2024). However, the investigation of judgment (and other cognitive) biases in the other 98% of species’ diversity, invertebrates, is greatly reduced. There is therefore a knowledge gap on the potential for affective states in invertebrates (Horváth *et al*., 2013). This is despite the behavioural complexity shown in invertebrate taxa such as insects.

Mental experience, emotion, and welfare in invertebrates have historically been under-considered and under-researched (Eisemann *et al*., 1984). Although recently interest and knowledge on related topics such as insect sentience has increased (Elwood *et al*., 2009; Elwood, 2011; Godfrey-Smith, 2021b), insight into affective states is limited (Horváth *et al*., 2013); restricted to eusocial species and those commonly used in laboratories (Perry & Baciadonna, 2017; Lucon-Xiccato *et al*., 2024), such as honeybees (Bateson *et al*., 2011; Schlüns *et al*., 2017), bumblebees (Solvi *et al*., 2016; Strang & Muth, 2023), ants (d’Ettorre *et al*., 2017; Wenig *et al*., 2022), and fruit flies (Deakin *et al*., 2018; Polizos *et al*., 2024). Understanding for all other insect orders aside from Diptera and Hymenoptera, and all hemimetabolous insects in general, is therefore absent. This is especially problematic if we consider how trillions of insects are slaughtered per year in the food and feed industry (Rowe, 2020; Klobučar & Fisher, 2023), plus many more exterminated in pest control and used in scientific research. If these animals possess affective states and can suffer, then we may need to consider adapting methods and legislation at a massive scale in order to limit the negative impact on the welfare of captive insects.

Honeybees (Bateson *et al*., 2011) and ants (Wenig *et al*., 2022) have always been popular groups in animal cognitive research. One group of insects that offers interesting cognitive capabilities for affective state research are cockroaches (Lucon-Xiccato *et al*., 2024). Cockroaches show emergent collective behaviour and even complex social cognition (Jeanson *et al*., 2005; Planas-Sitjà & Deneubourg, 2018; Planas-Sitjà *et al*., 2018; Günzel *et al*., 2020; Lu *et al*., 2023) and are used in the study of conditioning, learning and memory (Watanabe *et al*., 2003; Arican *et al*., 2020; Lu *et al*., 2023), and so are suitable for testing for cognitive biases. Further, several thousand tons of cockroaches are farmed annually (van Huis *et al*., 2024; Siddiqui *et al*., 2024) and destroyed as pests in uncounted numbers. However, we are unaware of reports of any cognitive bias test on any cockroach species. There is, therefore, an opportunity to perform a cognitive bias test on an understudied taxon of non-eusocial insects. Additionally, animal welfare science has historically been biased toward studying negative emotions, but it is now widely accepted that good animal welfare is not created from the absence of negative states but by the presence of positive ones (Boissy *et al*., 2007). Solvi *et al*. (2016) and Wenig *et al*. (2022) both induced positive judgement bias responses to ambiguous stimuli in bumblebees and ants, but their methods have been criticised (Baracchi *et al*., 2017). Both Solvi *et al*. (2016) and Wenig *et al*. (2022) use “unexpected” sucrose rewards as the method to induce the state of positive judgement bias to ambiguous stimuli, but the presence of that unexpected sucrose might be inducing a change in foraging motivation rather than inducing an optimistic affective state, increasing the individuals’ exploratory behaviours rather than influencing an emotion-like state (Baracchi *et al*., 2017).

Here, we developed a straightforward judgement bias test on *Blaptica dubia* cockroaches, following the parameters set out by Bateson *et al*. (2011) and Bethell (2015), allowing us to test for the presence of both pessimistic and optimistic affective states. We avoid previous issues with “unexpected” rewards by taking advantage of the fact that gregarious cockroaches are highly sensitive to the presence of other cockroaches when it comes to shelter choice (Günzel *et al*., 2020; Freeberg *et al*., 2024), spatial navigation (Lu *et al*., 2023) and even show collective group decision-making (Lihoreau *et al*., 2012). These social influences can be induced using the scents of other cockroaches of the same species (Günzel *et al*., 2020) or even different species (Freeberg *et al*., 2024). We therefore used the scents of opposite-sex conspecifics to induce a positive judgement bias in cockroaches instead of using ‘unexpected’ sucrose rewards.

First, to confirm cockroaches could be tested within a judgement bias paradigm, we tested whether we could condition cockroaches to associate one scent with reward (positive conditioned stimulus; CS+) and one scent with punishment (negative conditioned stimulus; CS-), and then recorded whether conditioned cockroaches would respond (by extending their maxilla-labia) to a range of ambiguous scents (mixtures of the two scents at different ratios). We predicted that cockroaches could be conditioned to associate a scent with a reward and that the strength of their response would depend on the ratio of the mixture. We then performed two experiments by comparing the responses between differently treated groups of cockroaches (e.g., those experiencing a negative or positive stimulus versus a control group). In the first experiment, we attempted to induce negative affective states by comparing cockroaches exposed to high light intensity (which is aversive to cockroaches, Szymanski, 1912, Yadav & Paul, 2024) to a control group kept at standard low light intensity. We predicted that cockroaches exposed to light would be less responsive to ambiguous stimuli than the control group, indicating negative affective states. In the second experiment, we attempted to induce positive affective states by exposing cockroaches to the scent of opposite-sex conspecifics, compared to a control group without this exposure. We predicted that the group exposed to cockroach scent would be more responsive to ambiguous stimuli, indicating positive affective states.

## Methodology

### Research subjects and harnessing

We used *B. dubia* cockroaches from a population kept at the University of Aberdeen since 2021. The cockroaches are typically kept at 28°C, 50% humidity, with a 12:12 light:dark cycle (lights on at 07:00, and so all experimental work is conducted in the photophase). They are provided egg cartons (cardboard) as shelters and are fed Sainsbury’s Complete Nutrition Adult Small Dog Dry Dog Food (for nutrition) and carrots (for hydration). We harnessed test subjects in custom-built harnessing devices dubbed ‘cockroach harnesses’ (see Fig. 1). The harness’s purpose is to allow freedom of movement for the subject’s head and antennae while restricting their lower body movement. We made the harnesses by cutting 30ml screw-top plastic vials to create plastic tubes with an interior diameter of approximately 22mm. We used thermoplastic sheets to shape plastic caps for the cut-off end of the tubes. We designed the plastic caps to have a gap (approximately 4mm wide) so that the plastic caps could be pinned between the head and the thorax. We placed the test subjects into the harnesses with tissue paper as extra support to ensure no movement was possible besides the head and the antennae; this also reduced the chances of the subjects escaping from the harnesses. Females were too broad and strong to be held in the harnesses without risk of escaping or being harmed by the harnessing process, and so we only used males in this study. We restricted the test subjects from food or water for 24 hours prior to being placed in harnesses. All test subjects were given 30 minutes to become accustomed to the harnesses while kept in a sheltered box (light intensity of 0.09 x10µmol m^2^/s).

**Figure 1.**
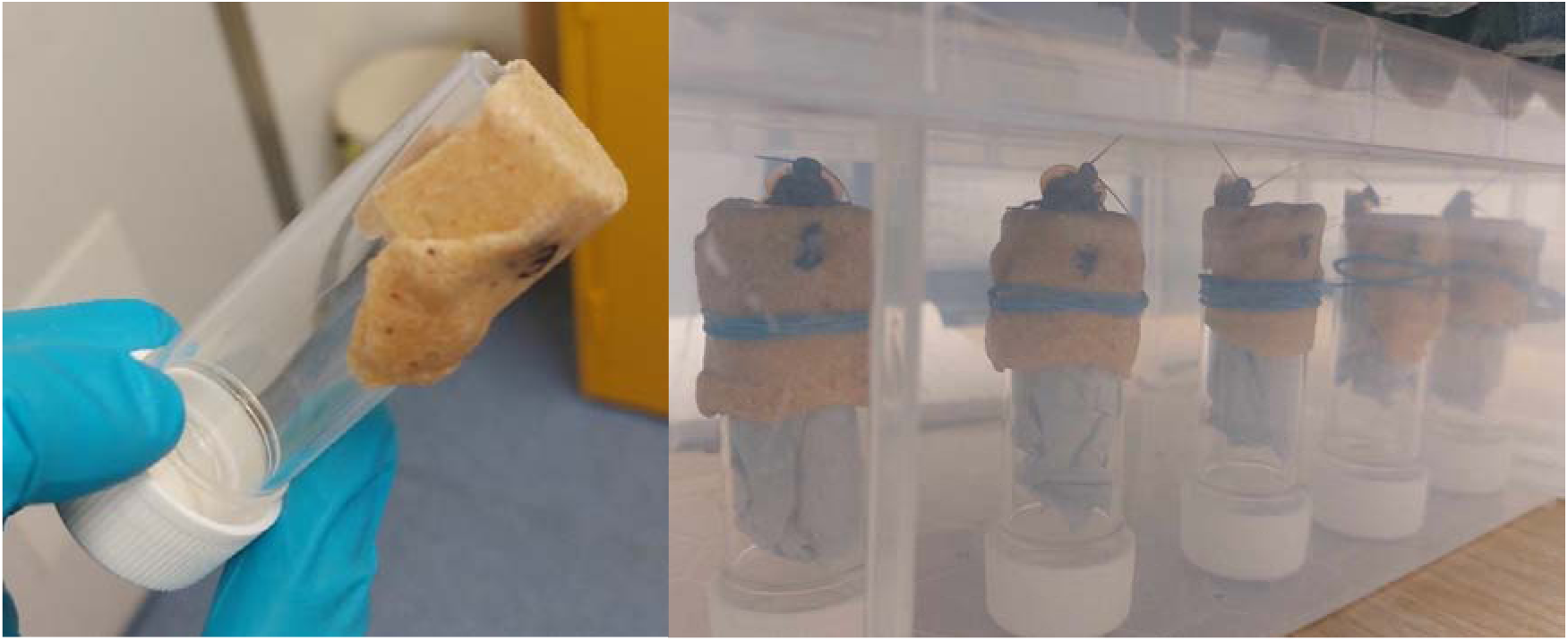
Cockroach harnesses. Left: An empty cockroach harness and Right: Cockroach harnesses in use during the experiment (pre-conditioning 30 minutes resting period).

### Conditioning procedure

We developed a conditioning procedure based on the approach used on honeybees from Bateson *et al*. (2011) and the methods used on *Periplaneta americana* cockroaches from Watanabe *et al*. (2003) and Arican *et al*. (2020). We subjected the cockroaches to 12 differential conditioning trials following a systematic order (ABABABABABAB; where A = CS+, B = CS-), with intertrial intervals of 5 minutes. Each trial involved the presenting of an odour for 3 seconds, followed by an appetitive (CS+) or aversive (CS-) outcome. We conditioned the cockroaches with two odours: isoamyl acetate (Mystic Moments; CAS: 123-92-2) and butyric acid (Mystic Moments; CAS: 107-92-6), following Arican *et al*. (2020). We presented odours by soaking 0.2ml of odour solution (99.50 to 99.70%) in cotton balls and held the cockroaches’ heads over a vial containing this ball for three seconds (Fig. 2). After presenting a subject with isoamyl acetate, we placed one drop of 1M sucrose solution on the mouth parts of the subject using a hypodermic syringe (CS+). Similarly, after we presented butyric acid we placed one drop of saltwater solution (CS-) with a different needle. We prepared the saltwater solution by mixing 6.48g of NaCl in 20 ml of demineralised water. When we presented the reward or punishment, we also included a cotton ball scented with the appropriate scent on skewered towards the base of the needle used to present the reward or punishment (Fig. 2).

**Figure 2.**
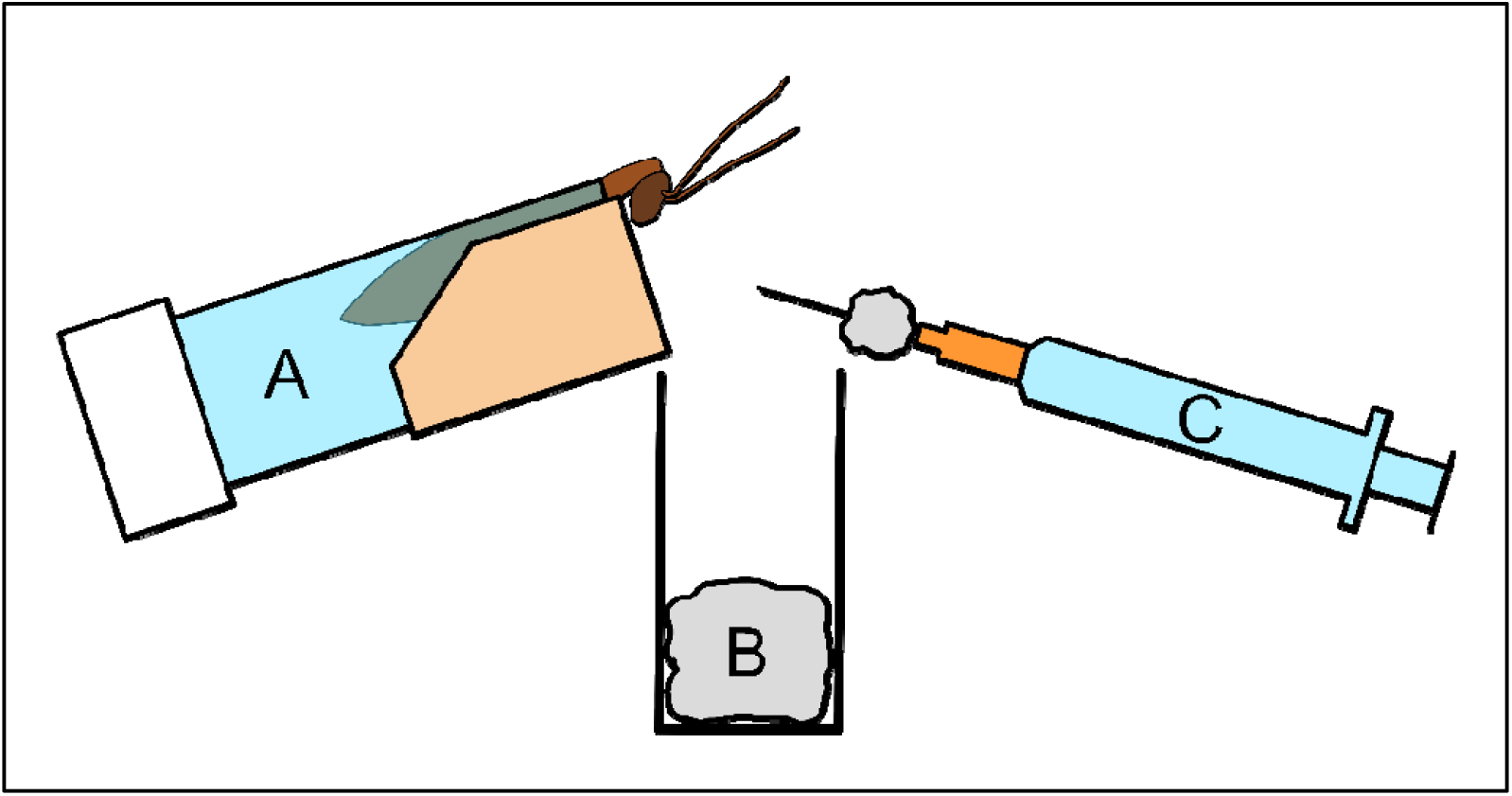
Cockroach conditioning set up. A: Cockroach harness with a cockroach. B: Glass vial containing scented cotton ball. C: Syringe filled with reward (sucrose solution) or punishment (saline solution) with scented cotton ball skewered on needle.

### Conditioning Effectiveness Tests

We measured conditioning effectiveness by whether the cockroach extended its mouthparts (maxilla-labia) toward the needle within 3 seconds (duration chosen following Arican *et al*. 2020) of being presented with an odour without any reinforcement outcomes (sucrose or saline solutions). Arican *et al*. (2020) refer to this response as the maxilla-labia response (MLR), an unconditioned response present in cockroaches when detecting the presence of food, a similar unconditioned response as the extension of the proboscis in honeybees (Bateson *et al*., 2011). To test the duration of the conditioning we repeated this presentation after 5, 30, and 60 minutes and 24 hours after the final conditioning trial. We measure the effectiveness of the conditioning by comparing the proportions of MLR responses (which are present/absent in a single trial) to Isoamyl acetate (CS+), Butyric acid (CS-), and unscented cotton balls (Control, C). In each test we presented cockroaches with the CS+, CS-, and C stimuli in a random order, and recorded their MLR responses. In between the 60 minute and 24-hour post-conditioning tests we removed the subjects from their harnesses and placed them into individual small boxes kept under shelter. We then strapped the subjects back into their harnesses 30 minutes before performing the post-24-hour conditioning effectiveness test, to become accustomed to the harnesses as before. We tested 30 males across all time points and another 14 males in just the 5-, 30-, and 60-minute time points. The tests occurred over 11 days.

### Negative Cognitive Bias Test

To test for negative cognitive biases, we created a range of ambiguous stimuli. We mixed the conditioning odours (isoamyl acetate and butyric acid) into five solutions of varying ratios with a total volume of 1ml: 9:1 (0.9ml of isoamyl acetate and 0.1ml of butyric acid), 7:3, 1:1, 3:7, and 1:9. First, following the procedure above, we conditioned a new set of cockroaches to respond positively to isoamyl acetate and not to butyric acid (we did not test for the effectiveness of the conditioning once the 12 conditioning trials were complete). After conditioning procedures, without removing subjects from their harnesses we kept half the cockroaches (the control group) under shelter (cardboard egg trays) for 1 hour, giving a light intensity of 0.04 x10µmol m^2^/s. The other half (negative stimulus group) we kept in an unsheltered transparent box for 1 hour, giving a light intensity of 0.88 x10µmol m m^2^/s. High light intensity is aversive and stressful to many cockroach species (Zhukovskaya et al., 2017; Yadav & Paul, 2024), including *B. dubia* (Freeberg & Fiset, 2023; Freeberg et al., 2024), as they are typically nocturnal (Bell et al. 2007; Laurent-Salazar et al. 2019). As such exposure to light is an effective aversive stimulus. After the hour, we presented the five mixed odour solutions to the subjects in a random order, presenting each odour for 5 seconds and recording the presence or absence of MLR. This is a slight increase over the 3 seconds used in the conditioning effectiveness tests described above, as we aimed to match the protocol of Bateson *et al*. (2011) as closely as possible for the cognitive bias testing. We tested cockroaches in batches of 5 cockroaches at a time, switching between sheltered and unsheltered each batch to reduce the effects of confounding variables, such as time of day. We assayed 35 individuals in each treatment group over a period of 12 days.

### Positive Cognitive Bias Test

Our test for the positive cognitive bias followed the same procedure as the negative bias test, except that the experimental treatment cockroaches were kept in a sheltered box alongside egg cartons removed from a stock box with adult female cockroaches (hence would be strongly scented of female cockroaches) for 1 hour, while the control group had new and so unscented egg cartons (both groups were also sheltered from light and experienced a light intensity of 0.01 x10µmol m^2^/s). Prior research attempting to induce positive affective states has used the provision of additional food but then confound detection by using a response to food as part of the test for affective state (Solvi *et al*. 2016; Baracchi *et al*., 2017; Wenig *et al*. 2022). We aim to avoid this by using a completely different stimulus type – the scent of potential mates. This should induce a positive affective state as this stimulus suggests a good environment to achieve fitness gains (Fraser & Duncan, 2023). For instance, housing canaries (*Serinus canaria*) in pairs compared to in isolation leads to a positive affective state (Lalot *et al*., 2017). We note here that this odour cue may induce enhanced attention to odours and/or increased risk-taking, which might affect responsiveness to the ambiguous cues in the absence of changes in affective states. We return to these points in the Discussion. However. these egg cartons were not present at the time of the test, so the male cockroaches could not show MLR *directly* due to the scent of females. We tested 19 individuals in each group over a period of 7 days, and so one batch only contained four males.

### Data analysis

We conducted all analyses in R (version 4.4.3) (R Core Team, 2025) and produced all graphs using the packages “dplyr” (Wickham *et al*., 2023) and “ggplot2” (Wickham, 2016).

We recorded the response data in a binary format (1 = response occurred, 0 = no response occurred), with repeated measures on individual cockroaches. We therefore used generalised linear mixed-effect models in the package “lme4” (Bates *et al*. 2015) for the conditioning effectiveness, negative bias, and positive bias experiments. In each model we set “nAGQ = 0” to allow convergence. We used a type II Analysis of Deviance in the package “car” (Fox & Weisberg, 2019) to generate p values. For the conditioning effectiveness test we included fixed effects of time (a categorical variable with levels of 5, 30, or 60 minutes or 24 hours) and condition (a categorical variable with levels of CS+, CS-, or C) and the interaction between these two terms, and a random effect of cockroach ID. If cockroaches were effectively conditioned, we would expect the main effect of condition to be significant, while if the strength of the conditioning changes over time we would also expect the interaction to be significant. We used the “emmeans” package (Lenth & Pisaskowski, 2026) to calculate the average (across all levels of time) log-odds ratios between each pair of C, CS+, and CS-.

For the negative and positive bias tests, we included the ratio of the two odours as a five-level categorical variable, the two-level categorical variable of treatment (control and either exposed to light or exposed to scent for the negative and positive bias tests respectively), and the interaction between odour ratio and treatment. Both models also included the random effect of cockroach ID. We expected that the cockroaches would be most responsive to the 9:1 ratio and continuously decrease in responsiveness to the 1:9 ratio. Additionally, if we successfully induced negative affective states, we expect the group exposed to light to be less responsive than the controls, while if we induced positive affect states the group exposed to the scent of conspecifics would be more responsive than controls. We used the emmeans package to calculate average log-odds ratios between control and exposed in each model, this time across all odour ratios.

We wanted to check whether it was reasonable to directly compare our two cognitive bias experiments. To do this we compared the proportion of cockroaches responding to each odour ratio in the two control groups between the two experiments with a Chi-squared test. Finding that the two control groups were not different would indicate the experiments could be reasonably compared.

## Results

### Conditioning Effectiveness

We found that cockroach responses to the different odour stimuli (CS+, CS-, C) were consistent across time points (interaction between time and condition: χ^2^_6_ = 0.553, p = 0.997). Responses to CS+ were always high and the responses to CS- or C always low (Fig. 3; main effect of condition: χ^2^_2_ = 36.048, p < 0.001, average log-odds ratio of C:CS- = 0.744, for C:CS+ = 6.60 ×10^−6^, and for CS-:CS+ = 8.90 ×10^−6^). The main effect of time was not significant (χ^2^_3_ = 0.950, p = 0.813). Cockroach ID had a variance of 1.687

**Figure 3.**
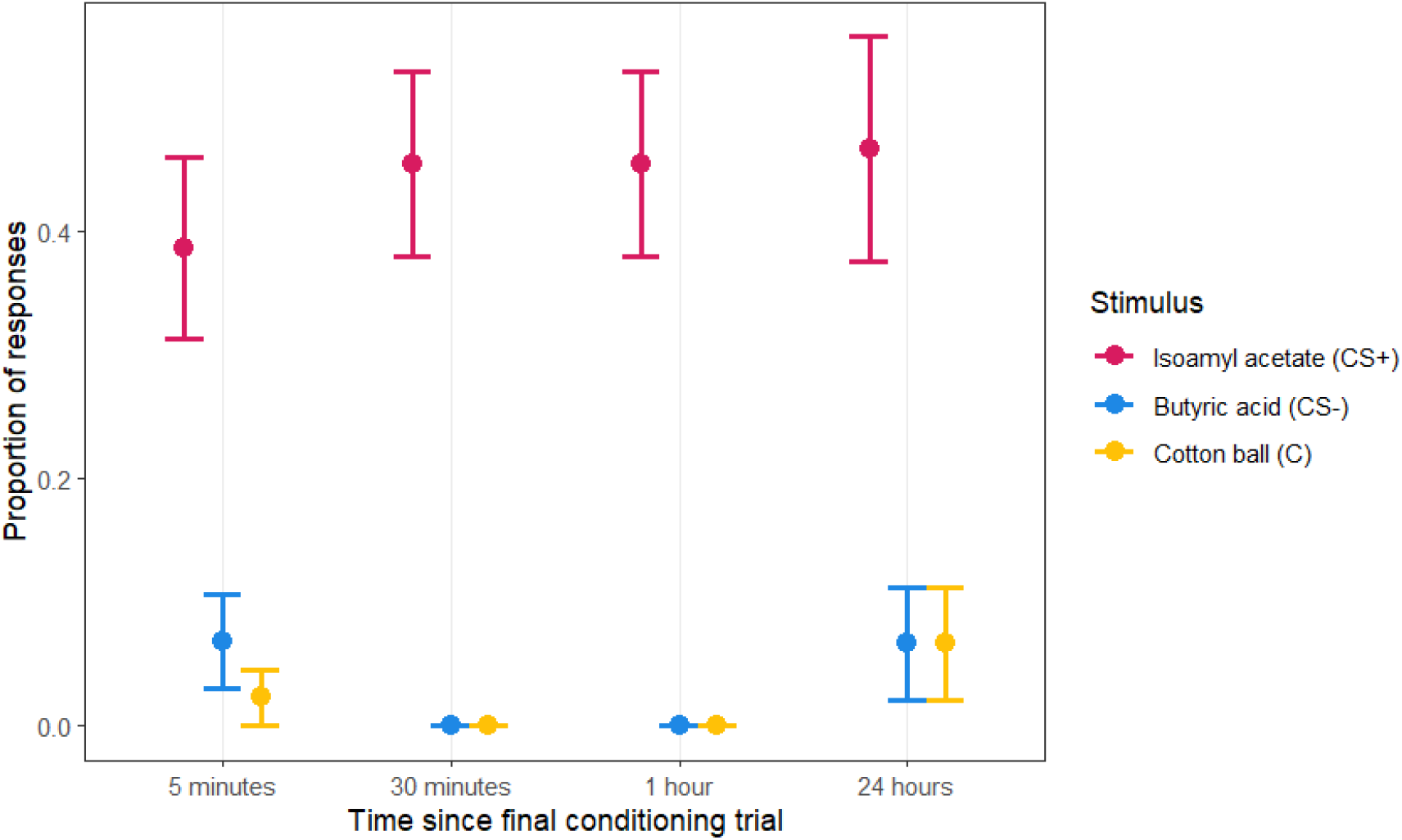
Proportion of cockroaches responding to unreinforced stimuli odours (Isoamyl acetate [CS+], Butyric acid [CS-], Cotton ball [C]) at different timepoints following conditioning. Points indicate proportion with error bars indicating standard error of the proportions.

### Pessimistic Cognitive Bias

We found that cockroaches were less responsive in the unsheltered condition (χ^2^_1_ = 3.794, p = 0.020, average log-odds ratio of sheltered:unsheltered = 5.67), indicating negative affective states. Responsiveness declined as the ratio of isoamyl acetate to butyric acid decreased (χ^2^_4_ = 85.4543, p < 0.001; Fig. 4). There was no interaction between treatment and ratio (χ^2^_4_ = 5.768, p = 0.217) indicating the effects of each variable were additive. The variance attributed to cockroach ID was 8.650.

**Figure 4.**
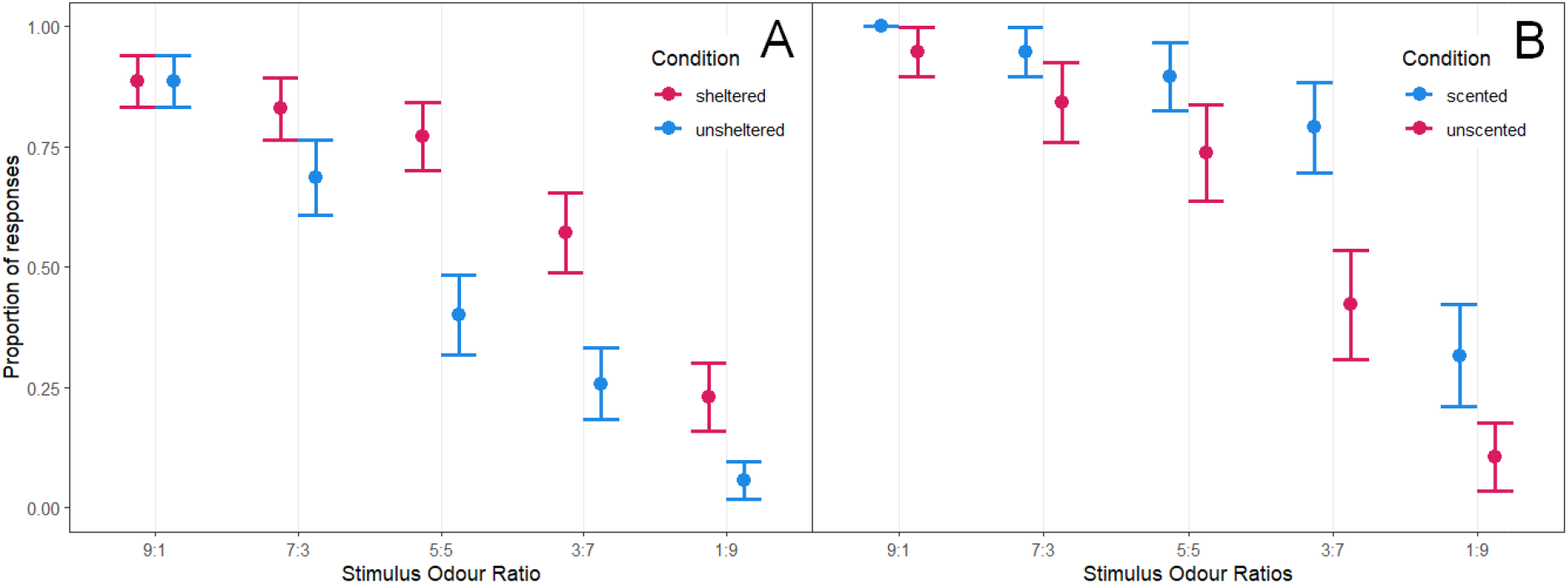
Cockroach responses to different ratios to isoamyl acetate to butyric acid by condition (A.: sheltered/unsheltered; B.: scented/unscented). Note that in both graphs the control group is shown in red and the treatment group in blue, but the direction of the effect of the treatment is opposite in the two experiments.

### Optimistic Cognitive Bias

We found that cockroaches were marginally more responsive in the scented condition (χ^2^_1_ = 3.794, p = 0.051, average log-odds ratio of scented:unscented = 178), providing tentative evidence consistent with positive affective states. Responsiveness declined as the ratio of isoamyl acetate to butyric acid decreased (χ^2^_4_ = 46.396, p < 0.001; Fig. 5). There was no interaction between treatment and ratio (χ^2^_4_ = 0.853, p = 0.931) demonstrating the effects of each variable were additive. Cockroach ID accounted for a variance of 8.131.

### Comparing controls

To confirm the two experiments were directly comparable we tested whether the response rates in the control conditions in both analyses (sheltered condition in the pessimistic cognitive bias test and unscented condition in the optimistic cognitive bias test) were different. We found that the two controls were similar (analysis of proportion of MLR responses: Chi-squared test, χ^2^_4_= 0.667, p = 0.955).

## Discussion

We were able to condition cockroaches to associate a scent with a food reward and another with punishment, and to respond to the former with an extension of their mouthparts. They then showed intermediate responses to mixtures of the two scents (ambiguous stimuli) and crucially were less responsive if they had experienced negative conditions (exposure to light) and (marginally) more responsive if they had previously experienced positive conditions (the scent of conspecifics). Together these results suggest *B. dubia* has the potential to possess affective states – the first such results in a cockroach and in fact any hemimetabolous insect.

Post-conditioned cockroaches showed good responsiveness (45%) towards the positive stimulus and limited to no response to the control or negative stimuli. These response rates indicated a significant enough difference to justify judgement bias testing. The 40-45% overall response rate to isoamyl acetate is slightly lower response rate than that seen in Arican *et al*. (2020), which showed a 50-60% response rate to isoamyl acetate when conditioning *Periplaneta americana*. The difference in post-conditioning response rates between the species might suggest variability of conditioning rates within the taxon, which has not been explored in cockroaches before. However, we also showed a response rate of 88% in the 9:1 ratio for the negative cognitive bias tests, nearly double the rate we found in the conditioning tests. The difference in response rate might be due to the difference in time given to respond for the conditioning effectiveness tests and cognitive bias tests. The conditioning effectiveness had a 3-second response recording time, following the protocol set out by Arican *et al*. (2020), while cognitive bias tests had a 5-second response recording time, following the protocol by Bateson *et al*. (2011), which would allow a higher response rate.

The cockroaches we exposed to a negative stimulus showed a significant pessimistic bias toward the range of ambiguous scent ratios. Responses to the 1:1 ratio showed the most prominent difference of 40% (unsheltered) and 77% (sheltered [control]). This is a comparable, if not larger, difference to what Bateson *et al*. (2011) found for honeybees, which showed a difference of approximately 45% (shaken) and 60% (non-shaken [control]). Therefore, both cockroaches and honeybees can be induced to show pessimistic cognitive biases at similar levels. Most other insect affective state research to date has been based on eusocial species (ants, honeybees, bumblebees); our study on a non-eusocial gregarious insect species offers up the opportunity to compare cognitive biases between eusocial and non-eusocial insects (Lihoreau *et al*., 2012; Friedman *et al*., 2020; Couzin-Fuchs & Ayali, 2021). Gregarious cockroach species in particular represent a comparative model for understanding the evolution of insect sociality (Lihoreau *et al*., 2012), which might have important consequences for their cognitive ability. We suggest expanding comparable cognitive bias testing across *Blattodea*, focusing on comparing cognitive bias tests among solitary, gregarious, family-living, and eusocial cockroaches (termites).

The cockroaches in the optimistic judgement bias experiment were typically more responsive to ambiguous scents after exposure to cockroach-scented cardboard, although the result was marginally non-significant. This supports previous work by Solvi *et al*. (2016) (bumblebees) and Wenig *et al*. (2022) (ants), suggesting that positive affective states in insects can be induced. Further, our results provide tentative evidence for the presence of an induced optimistic bias state without using an ‘unexpected’ sucrose reward. It has been suggested that the ‘unexpected’ sucrose reward used to induce positive affective states by Solvi *et al*. (2016) acts as a motivational trigger for increased exploratory behaviour, and that increased exploratory behaviour is then recorded in the judgement bias test as optimistic responses (Baracchi *et al*., 2017). Since we used cockroach scent to induce the optimistic state rather than ‘surprise’ sucrose rewards, which were then absent when testing, the MLR should be directly in response to the odour we presented, giving a stronger test for positive affective states. Cognitive bias tests done on vertebrates also use various methods of inducing optimistic affective states without the use of ‘unexpected’ food rewards (Clegg, 2018), such as positive human interaction (Brajon *et al*., 2015), environmental complexity (Zidar *et al*., 2018), and release from constraints (Doyle *et al*., 2010), and so future work on insects could explore these to find the most reliable and precise. We note here that, as we used the scent of females to induce positive affective states in males, it is possible that the marginally increased MLR might be because of increased mating-related motivation, which might increase the risk-taking behaviour in terms of responding to ambiguous stimuli (although the testing odours were presented separately to the scent of females). However, mate-finding is initiated by females in Blattaria cockroach species (Lizée et al., 2017), a grouping including *B. dubia*, which might limit how much we expect risk-taking behaviour to be increased in males. Nevertheless, as we are in the early stages of positive affective state research in insects, and as our approach to inducing a supposed positive state is novel, the aspect of our results on positive cognitive bias should be interpreted cautiously.

Our results suggesting that the presence of cockroach scent can induce behaviours consistent with an optimistic affective state implies that a process that *looks like* “emotional contagion” could be possible in cockroaches. Emotional contagions are the mirroring and synchronising of moods and emotions between two or more individuals (Špinka, 2012; Herrando & Constantinides, 2021). For example, play calls in kea parrot (*Nestor notabilis*) elicit higher rates of play behaviour in birds hearing the calls (Schwing *et al*., 2017). In the case of cockroaches, their decisions are affected by both the tactile presence of other cockroaches (Lihoreau & Rivault, 2008; Lihoreau *et al*., 2010) and scents (Lihoreau *et al*., 2010; Günzel *et al*., 2020). These results combined with ours raises the possibility communication between cockroaches based on scent or tactile cues could lead to them possessing a similar affective state to other cockroaches they contact. Pleasant or unpleasant experiences could alter chemical cues, which could then be picked up by other cockroaches who did not experience those experiences, altering their affective states (which looks identical to emotional contagion, even if the mechanisms in other taxa may differ). Wenig *et al*. (2022) were unable to find any signs of emotional contagions within black garden ants (*Lagus niger*), as social information, such as trails or pheromones did not affect judgment bias. However, Romero-González *et al*. (2025) have recently shown that bumble bees (*Bombus terrestris*) induced into a positive affective state through an unexpected sucrose reward transmitted this state to another bee. Further, this transfer relied on visual information and could occur when bees were separated by an acrylic sheet but not when they interacted in the dark (Romero-González *et al*., 2025). Since we specifically tested for an impact of some social information transmitted by scent on judgement bias, we would expect emotional contagions to happen through this medium (or tactile cues) in cockroaches instead of via visual cues. While emotional contagions are thought to be widespread amongst social vertebrates (Düpjan *et al*., 2020; Pérez-Manrique & Gomila, 2021), evidence more widely is limited. Designing assays to test for emotional contagion in a wider range of invertebrates that are comparable to the highest-standard tests in vertebrates would allow us to assess the spread of this phenomenon across taxa.

Cockroaches showing affective states has further implications for the concept of mental experience and emotion in invertebrate animals. While in the UK, the recognition of consciousness and sentience within invertebrates has been restricted to decapods and cephalopods (DEFRA, 2021), the expansion of evidence of mental experience across various groups of insects and the increased public and academic interest in invertebrate consciousness and welfare (Drinkwater *et al*., 2019; Godfrey-Smith, 2021a; de Waal & Andrews, 2022) might warrant another expansion of sentient categorisation (Perry & Baciadonna, 2017). The presence of pessimistic and, possibly, optimistic states in cockroaches also brings into question how they are kept in captivity. The presence of affective states necessitates their consideration for the welfare of captive animals (zoo or livestock), by making changes to amplify positive affective states with enrichment and optimum conditions and minimise negative affect states by minimising pain, fear, and stress (Clegg, 2018; Düpjan *et al*., 2020). With increased interest in honey production (Phiri *et al*., 2022) and insect production for food and feed (Lambert, *et al*., 2021; Klobučar & Fisher, 2023), welfare consideration for invertebrates is a logical extension for existing practice.

To conclude, our results are consistent with the notion that cockroaches possess affective states. These findings support presence of emotion like states in insects (both eusocial and non) and expand the presence of cognitive bias to unresearched taxa (Perry & Baciadonna, 2017).

## Acknowledgements

We thank Keith Lockhart for his hard work maintaining the cockroach stock population and general advice. We also thank Ryan Heffernan and Ben Saber-Sheikh for constructive discussions and Esmé Holme, Nicole McCue, and two anonymous reviewers for reviewing an earlier draft.

